# Wheel-running exercise before and during gestation against acute and sensitized cocaine psychomotor-activation in offspring

**DOI:** 10.1101/477505

**Authors:** Louis-Ferdinand Lespine, Alain Plenevaux, Ezio Tirelli

**Affiliations:** Université de Liège, Département de Psychologie, Quartier Agora - Place des orateurs, 2 (B32), 4000 Liège, Belgique; Université de Liège, Département de Chimie, Quartier Agora - Allée du 6 Août, 8 (B30), 4000 Liège, Belgique

**Keywords:** Wheel-running, gestation, prenatal, mice, offspring, cocaine sensitization

## Abstract

While animal research has consistently reported preventive effects of exercise against drug abuse vulnerability, little is known about the influence of the developmental stage during which exercise is displayed on addictive drugs responsiveness. This study aimed to determine whether prenatal exercise could attenuate acute cocaine reactivity and psychomotor sensitization in youth offspring. We used a split-plot factorial design where C57BL/6J females were randomly assigned into sedentary or exercised (wheel-running) conditions before and during gestation, the wheels being removed on gestational day 18. Offspring were weaned, gendered and individually housed on 24-28 days old. At 38-42 days old, they were tested for their acute psychomotor responsiveness to 8 mg/kg cocaine and their initiation of sensitization over 8 additional once-daily administrations, the long-term expression of sensitization occurring 30 days later. Adolescent females born from exercised mothers were much less responsive to the acute psychomotor-stimulating effect of cocaine than those born from sedentary mothers (*d* = 0.75, *p* = .02), whereas there was no evidence for such a difference in males (*d* = 0.34, *p* = .17). However, we did not find sizeable attenuating effects of prenatal exercise on the initiation and the long-term expression of the psychomotor-activating effect of cocaine, in either sex (Cohen’s *d*s varying from −0.23 to 0.39). These results suggest that prenatal exercise may induce initial protection against cocaine responsiveness in youth females, a finding that warrants further research.

## 1 Introduction

Epidemiological studies reported a negative association between physical exercise or sports participation and the initiation of drugs of abuse consumption (e.g., tobacco and cannabis smoking) [1,2]. Likewise, preclinical research has shown that exercised rodents through wheel-running - the most popular model of aerobic exercise in this field -, exhibit attenuated rates of acquisition or motivation of self-administration of various addictive drugs including cocaine, compared to their unexercised counterparts [2,3]. Consistent with self-administration reports, wheel-running exercise has also been shown to be effective at reducing the acute and chronic locomotor-stimulating effects of cocaine as well as the expression of sensitization to those effects [4-7], a phenomenon involved in addictive processes [8,9].

Recently, we observed that wheel-running exercise during adolescence (from 28 to 50 days of age), but not adulthood (from 77 to 99 days of age), induced long-term protection against cocaine psychomotor reactivity in females and males tested later in life [7]. These results suggest that early-life periods may be particularly sensitive to the (long-term) benefits of exercise, presumably because of, at least in part, structural and functional brain plasticity in regions involved in motivation and reward that continue to mature during adolescence [10,11].

We believe that the understanding of the interaction between exercise and development would deserve further investigations. In particular, it is well-known that some environmental stimuli during prenatal life (before or during gestation, or both) can subsequently affect the development and phenotypes of progeny.

In the absence of obstetric complications or medical contraindications, pregnant women are encouraged to participate in aerobic activities compatible with their exercise history [12,13]. Exercise during pregnancy is associated with health benefits regarding both maternal and pregnancy outcomes without evidence for detrimental effects on the fetus and neonate [13-15]. Although limited, a few studies have also reported a positive association between prenatal exercise and neurodevelopmental measures in offspring [16].

Preclinical research has shown that treadmill or wheel-running exercise during gestation (the exercise regimen sometimes starting before conception), can induce positive effects on several biological and behavioural endpoints in offspring, in particular, pro-cognitive effects [17-20]. For example, Robinson and Bucci [20] have shown that rats born from exercised mothers during pregnancy and tested in adulthood (from 60 days of age), exhibited better capabilities of discriminating between novel and familiar objects compared to rats born from sedentary ones. In spite of these encouraging evidence, we are unaware of experiments investigating the effects of prenatal exercise on drugs of abuse susceptibility in offspring. The present study attempted to determine the extent to which voluntary exercise performed before and during gestation could affect the initiation and the expression of psychomotor sensitization induced by a rewarding dose of cocaine in young female and male offspring. More specifically, we assessed cocaine-induced acute locomotor reactivity, locomotor sensitization developing over 9 once-daily sessions, and the long-term expression of the sensitized response in C57BL/6J mice born from mothers housed either with or without a running wheel before and during gestation.

## 2 Material and methods

The protocol was made available on the Open Repository and Bibliography of the University of Liège before the beginning of experimental testing (http://hdl.handle.net/2268/207692).

### 2.1 Ethical aspects

All experimental treatments and animal maintenance were reviewed by the University of Liège Animal Care and Experimentation Committee (animal subjects review board), which gave its approval according to the Belgian implementation of the animal welfare guidelines laid down by the European Union (“Arrêté Royal relatif à la protection des animaux d’expérience” released on 23 May 2013, and “Directive 2010/63/EU of the European Parliament and of the Council of 22 September 2010 on the protection of animals used for scientific purposes”). All efforts were made to minimize the number of animals used and their suffering. Moreover, the ARRIVE guidelines (Animal Research Reporting In Vivo Experiments), which have been developed to improve the quality of experimenting and reporting in animals studies, were followed as closely as possible [21].

### 2.2 Breeders housing

A total of 168 females and 84 males C57BL/6J mice were used for breeding and obtained at 6 weeks old from JANVIER, Le-Genest-Saint-Isle, France. Although these quantities were slightly below than those anticipated and mentioned in our protocol (N=192 females and N=96), this “deviation” has had no impact on our hypotheses, sample size (offspring), testing procedure or the statistical analysis plan. The choice of C57BL/6J strain was based on its extensive use in addiction research and previous exercise-related experiments performed in our lab [5-7]. Upon arrival, mice were housed in groups of eight (same-sex) in sizeable transparent polycarbonate cages (38.2 × 22 cm surface × 15 cm height; TECHNIPLAST, Milano, Italy) for one week of acclimation. On the following day, all mice were individually housed in smaller TECHNIPLAST transparent polycarbonate cages (32.5 × 17 cm surface × 14 cm height) with pine sawdust bedding, between-animal visual, olfactory and acoustic interactions remaining possible. Tap water and food (standard pellets, CARFIL QUALITY, Oud-Turnhout, Belgium) were continuously available. The animal room was maintained on a 12:12 h light-dark cycle (lights on at 07.00 a.m.), at an ambient temperature of 20-23°C. For females, the two housing conditions were defined by the presence or the absence of a running wheel on the surface of the tub. A running wheel was made of an orange polycarbonate saucer-shaped disk (diameter 15 cm, circumference 37.8 cm; allowing an open running surface) mounted on a plastic cup-shaped base (height 4.5 cm) via a bearing pin so as to being inclined from the vertical plane at an angle of 35° (ENV-044, Med Associates; St Albans, VT, USA). The base was fixed on a stable, transparent acryl-glass plate. Running exercise was monitored continuously using a wireless system, each wheel being connected to a USB interface hub (DIG-804, Med Associates) which relayed data to a Wheel Manager Software (SOF-860, Med Associates).

### 2.3 Experimental design and procedure

Experimental timeline and general design are presented in Figure 1. Based on a computer-generated randomization schedule, females were randomly assigned to exercise (EX, n=84) or sedentary (SED, n=84) housing. After 20 days in these conditions, they were housed with a breeder male for mating (two females for one male) during a 12-to-60 hour-period depending on the presence of a vaginal plug checked every morning. Then, females returned in the same pre-mating conditions (EX or SED), and the day was designated as gestational day 0 (GD-0). A total of 71 females (42.3%) showed gestation (EX, n=32; SED n=39). Wheels were removed from cages on GD-18 to minimize the risk of negatively influencing parturition. Mice were checked for birth twice daily.

**Figure 1.**
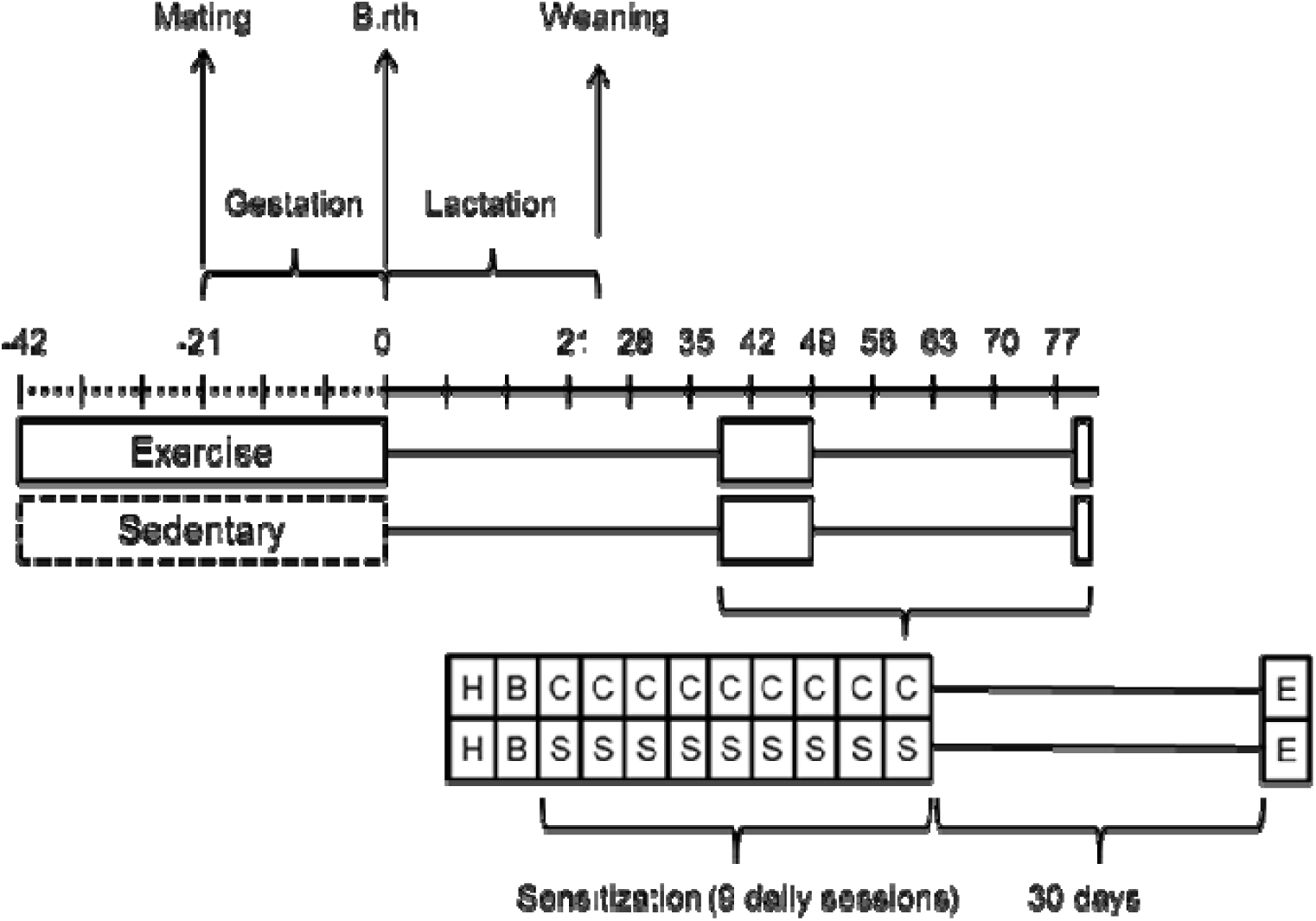
Experimental timeline and design. Seven-week-old females were housed individually either in the presence (EX) or the absence (SED) of a running wheel for 3 weeks followed by a 12-to-60-hour period of mating (two females for one male). Females returned into their respective cages (in the same pre-mating conditions, either EX or SED) and the day was designated as gestational day 0 (GD0). Wheels were removed on GD18. Day of birth was designated as postnatal day 0 (PND0), and pups were left undisturbed together with their mothers during the lactation period. Offspring were weaned on 24-28 days old, gendered, and individually housed. Testing began on 38-42 days old. The experiment comprised eight experimental groups incorporated in a split-plot factorial design with N=136, n=17; females and males (2 levels) from each prenatal housing condition (2 levels) receiving either cocaine or saline (2 levels). H: habituation session (to familiarize animals to the novelty of the test context without neither injection nor measures); B: baseline session, the 2nd once-daily session assessing the baseline activity under saline; C: cocaine intraperitoneal administration (9 once-daily sessions); S: saline administration (9 once-daily sessions; control animals). E: session on which the expression of the sensitization was assessed 30 days after the last sensitizing injection, under the previous pharmacological treatments.

The day of birth was designated as postnatal day 0 (PND-0), and pups were left undisturbed together with their mothers during the lactation period. At birth, 9 litters were excluded due to death or not viable neonates (EX, n=4; SED, n=5).

As the number of pups in C57BL/6J average 6, and because we wanted each experimental group (four groups; females and males receiving cocaine or saline) within each housing condition (EX or SED) to be represented in each litter by one mouse, only litters that comprised (at least) two females and two males were included. To employ the most homogeneous litters as possible, we also excluded litters with a number of pups ≥ 8. A total of 38 litters met these criteria (EX, n=17; SED, n=21). To obtain an equal sample size, 4 sedentary litters were excluded (randomly). Therefore, 34 litters were included in this experiment (EX, n=17; SED, n=17). Dams and pups that were not involved in this study were kept for either animal husbandry or the development of future experiments.

Since the four possible treatments (female/cocaine, female/saline, male/cocaine, male/saline) were represented within each litter with one mouse, litter was incorporated as a random factor in a split-plot factorial design with the housing condition (2 levels: EX, SED) as a between-block factor and sex (2 levels: female, male) and drug (2 levels: cocaine, saline) as within-block factors [23] with a total of eight experimental groups with n=17 (as independent experimental units). Supplement 1 presents the flowchart of dams/litters and the split-plot factorial design. This sample size was based on our previous experiment [7] where a 3-week regimen of exercise during adolescence induced a long-term (post-exercise) protective effect against cocaine responsiveness in females and males C57BL/6J, the smallest effect being observed in males (interaction between housing condition and drug found at □^2^p = 0.07 or *d* = 0.53 based on crossed contrasts; see “data analysis” section 2.6).

The eight experimental groups were as follows. (1) EX/Female/Cocaine: female born from mothers exercised before and during gestation and receiving 8 mg/kg cocaine during testing. (2) EX/Female/Saline group: female born from mothers exercised before and during gestation and receiving saline during testing. (3) EX/Male/Cocaine group: male born from mothers exercised before and during gestation and receiving 8 mg/kg cocaine during testing. (4) EX/Male/Saline group: male born from mothers exercised before and during gestation and receiving saline during testing. (5) SED/Female/Cocaine group: female born from mothers unexercised before and during gestation and receiving 8 mg/kg cocaine during testing. (6) SED/Female/Saline group: female born from mothers unexercised before and during gestation and receiving saline during testing. (7) SED/Male/Cocaine group: male born from mothers unexercised before and during gestation and receiving 8 mg/kg cocaine during testing. (8) SED/Male/Saline group: male born from mothers unexercised before and during gestation and receiving saline during testing.

Offspring were weaned on 24-28 days old, gendered, and individually housed in polycarbonate cages (30 × 12 cm surface × 13 cm height) with pine sawdust bedding, tap water and food (standard pellets, CARFIL QUALITY, Oud-Turnhout, Belgium) being continuously available. Behavioural testing began on 38-42 days old (adolescence [22]) and was identical to our previous study [7], comprising the following four phases. (1) A 1^st^ session to familiarize animals with the novelty of the test context without neither injection nor measures. (2) A 2^nd^ drug-free session evaluating the baseline psychomotor activity under saline. (3) Nine once-daily injections of cocaine or saline, initiating psychomotor sensitization after the baseline session. (4) Taking place 30 days after the 9^th^ (last) cocaine injection, a single-test assessing the long-term expression of sensitization on which animals received their previous respective pharmacological treatment. Three days before the testing began, all mice were familiarized with handling through saline injection in the animal room. Throughout testing, mice were weighed and received their pharmacological treatment right before being placed in the test chamber, the recording of ambulatory crossings lasting 30 min in all sessions. Experimental blinding was not realized because the unique experimenter inevitably knew the housing condition and the pharmacological treatment of each mouse.

### 2.4 Drug treatments

(−)-Cocaine hydrochloride (BELGOPIA, Louvain-La-Neuve, Belgium), dissolved in an isotonic saline solution (0.9% NaCl), was injected at a dose of 8 mg/kg in a volume of 0.01 ml/g of body weight, the control treatment consisting of an equal volume of isotonic saline solution. All injections were given intraperitoneally (i.p.). The dose and route of administration were based on our previous studies [6,7].

### 2.5 Behavioural test chambers

A battery of eight home-made chambers, connected to a custom written software for data collection, was used to measure mice psychomotor activity. Each activity chamber was constituted of a removable transparent polycarbonate tub (22 × 12 cm surface × 12 cm height), embedded onto a black-paint wooden plank serving as a stable base. The lid was made of a transparent perforated acryl-glass tablet. Two photocell sources and detectors were mounted on the plank such that infrared light-beams were located on the two long sides of the tub at 2-cm heights from the floor, 8-cm apart and spaced 6.5 cm from each end of the tub. Psychomotor activity (locomotion) was measured in terms of crossings detected by the beams, one crossing count being recorded every time an ambulating mouse broke the two parallel beams successively. The activity chambers were individually encased in sound-attenuated shells that were artificially ventilated and illuminated by a white light bulb during testing. Each shell door comprised a window allowing periodic surveillance.

### 2.6 Data analysis

Inferential statistics were computed on the following data. (1) The acute responsiveness to the psychomotor-activating effects of cocaine scored as the absolute difference between the values derived from the first cocaine session and those of the baseline session. (2) The overall responsiveness to the psychomotor-activating effects of cocaine over the initiation of sensitization (9 sessions) scored as the area under the curve with respect to zero. (3) The sensitization increment (development) of the psychomotor-activating effects of cocaine over the initiation of sensitization scored as the area under the curve with respect to the first cocaine session (AUC increase; calculation formulas are detailed in Pruessner and colleagues [24]). (4) The psychomotor activity exhibited during the expression of sensitization.

Each set of data was first analyzed by a split-plot factorial ANOVA in which the housing condition (2 levels: EX *vs*. SED) was incorporated as a between-block factor, with the sex (2 levels: female *vs*. male) and the drug treatment (2 levels: cocaine *vs*. saline) as within-block factors [23], estimation of the effect sizes being given by partial eta-squared (□^2^p), classifiable as small (0.01), medium (0.06) or large (0.14) [25].

The primary outcomes, defined by our *a priori* hypotheses, were given by planned crossed contrasts testing interactions of interest (see [26]). Specifically, our crossed contrasts, separately computed in females and males, compared the difference (or the effect) between the values of the EX/COC and EX/SAL groups with the difference between the values of the SED/COC and SED/SAL groups. Each contrast was derived from the appropriate mean-square error term (MSE) provided by the ANOVA, the statistical significance threshold being set at 0.025 (Bonferroni correction). One-tailed *t*-tests were used for these comparisons since mice from SED mothers were expected to display greater cocaine responsiveness than those from EX mothers. Effect sizes were estimated and given by Cohen’s *d*s (calculated from the corresponding *t*s and dfs), classifiable as very small (0.01), small (0.20), medium (0.50), large (0.80), very large (1.2) and huge (2.0) [27].

As secondary outcomes, the basic psychomotor-activating effects of cocaine within each group was ascertained using one-tailed *t*-tests (cocaine *vs*. saline) taken at a statistical significance threshold of 0.05, adjustments being unnecessary due to oversized effects. The graphs present means ± 95% confidence intervals (CI).

## 3 Results

### 3.1 Prenatal wheel-running activity

As shown in Figure 2, mice housed with a wheel showed a typical increase in running activity (expressed in kilometers / 12h) over two weeks, the plateau being reached from around the 10^th^ day (pre-mating period). A drop in running distances was observed immediately after mating (GD0), although mice maintained their running activity for a few days before progressively decreasing it as their pregnancy went forward. This pattern has been previously observed elsewhere [28-30] and in our lab (Supplement 2, Panel B). This phenomenon is unlikely to result from the forced encounter with the sire since non-gravid (non-pregnant) females did not exhibit it (Supplement 2, Panels A and B). Therefore, one can reasonably suggest that this sharp decrease was mainly due to their pregnancy-related physiological state.

**Figure 2.**
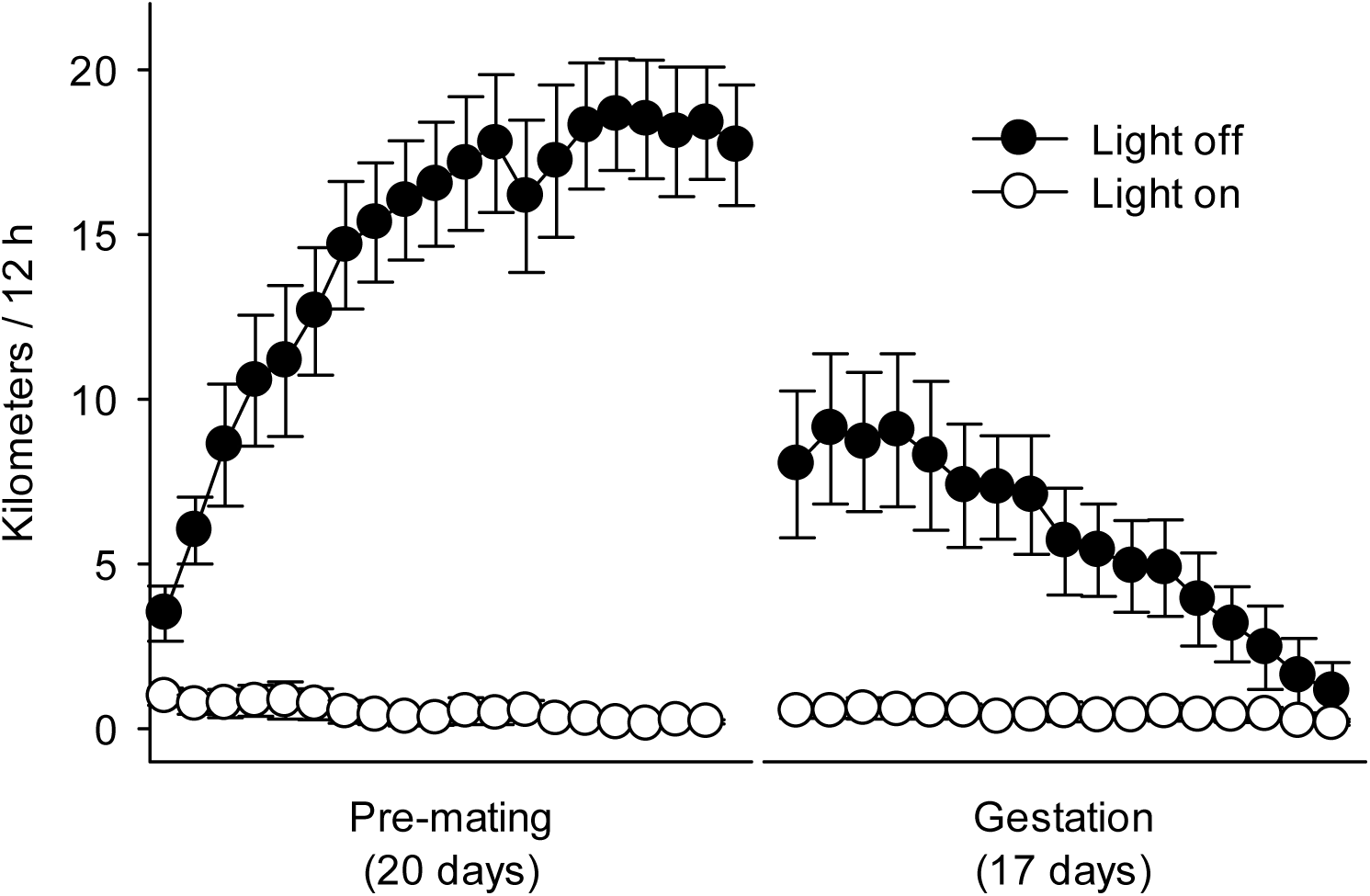
Wheel-running activity. Nocturnal (light off) and diurnal (light on) wheel-running activity (in kilometers / 12h) before and during gestation in gravid females used in this experiment (n=17). No inferential statistics were conducted on these data. Bars represent 95% CI.

### 3.2 Acute psychomotor responsiveness

Figure 3 depicts the acute responsiveness to the psychomotor-activating effects of cocaine in females (panels A and B) and males (panels C and D) born from mothers housed either with (EX) or without (SED) a running wheel before and during gestation. Panels B and D describe the time-course of the responsiveness over the six 5-min within-session intervals (no inferential statistics were computed on these data). The split-plot ANOVA resulted in a strong significant housing-by-drug interaction (□^2^p = .15, F_(1,32)_ = 5.75, *p* = .02) along with a negligible non-significant housing-by-drug-by-sex interaction (□^2^p = .01, F_(1,32)_ = 0.36, *p* = .55). Importantly, planned crossed contrasts indicated that the attenuating effect of prenatal exercise was significant and strong in females (*d* = 0.75, *t*_(32)_ = 2.11 at *p* = .02), but modest and non-significant in males (*d* = 0.34, *t*_(32)_ = 0.96 at *p* = .17). Statistics related to the basic psychomotor-activating effects of cocaine within each experimental group are provided in Table 2.

**Figure 3.**
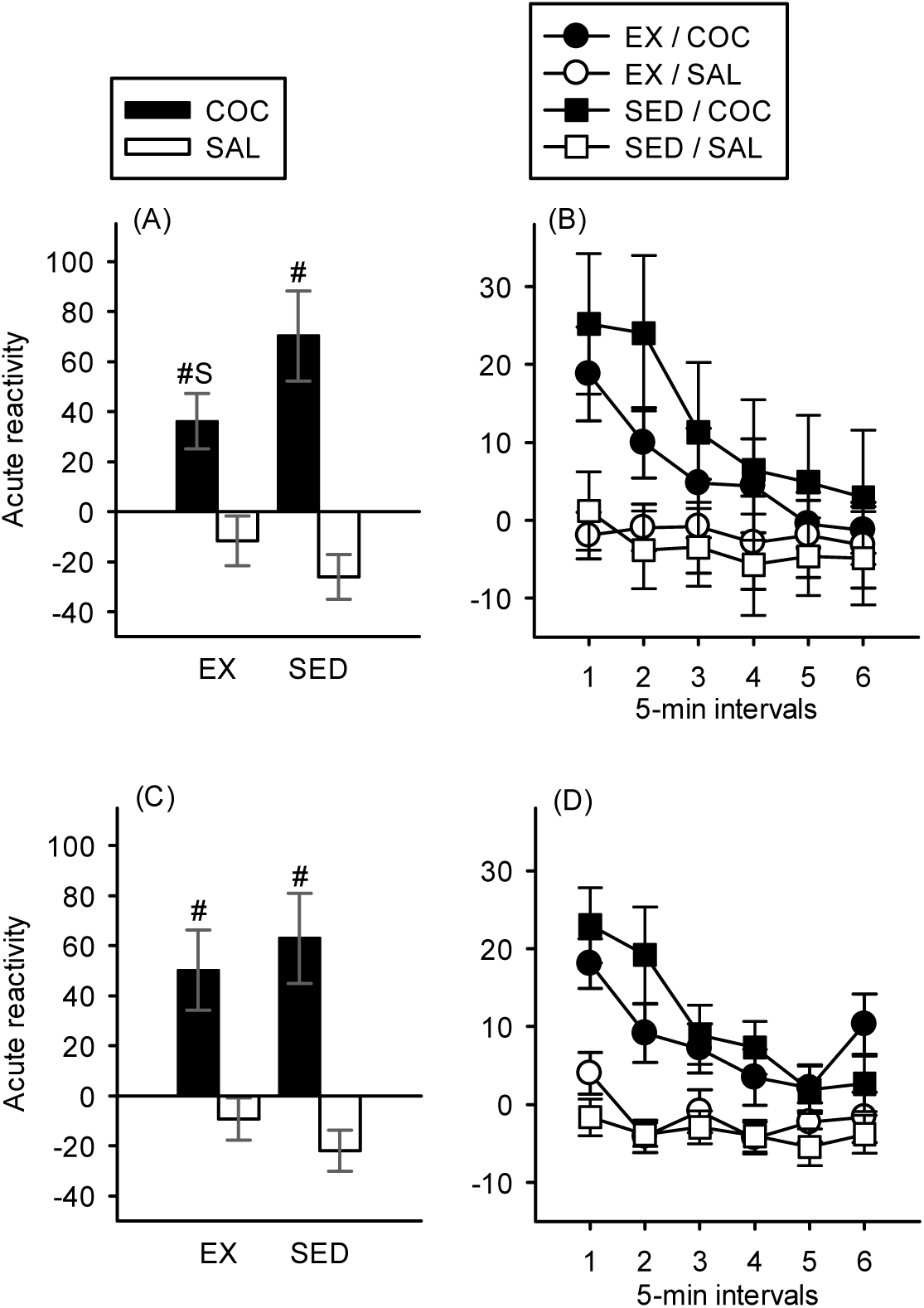
Acute reactivity. Left panels present acute psychomotor responsiveness to cocaine in females (A) and males (C) born from exercised (EX) or sedentary (SED) mice before and during gestation and tested at 38-42 days of age. Acute psychomotor responsiveness was scored as the absolute difference between values from the first cocaine session and the baseline session. S: significant interaction-related difference between the effects displayed by females born from exercised mothers (EX/COC *vs*. EX/SAL groups) and that measured in females born from sedentary mice (SED/COC *vs*. SED/SAL) taken at a *p*-level of 0.025. #: significant higher level than the corresponding saline group taken at a *p*-level of 0.05. Right panels describe time-course of acute psychomotor reactivity over the six 5-min within-session intervals in females (B) and males (D), represented by the absolute difference between values from the first cocaine session and the baseline session. No inferential statistics were conducted on these data. Bars represent 95% CI.

### 3.3 Initiation and expression of psychomotor sensitization

As depicted in Figure 4, there was no evidence for a significant difference in cocaine responsiveness between mice born from exercised mothers and those born from sedentary animals during the prenatal period, whichever experimental outcome was considered (Cohen’s *d*s varying from −0.23 to 0.39). Table 1 summarizes the main statistics related to the crossed contrasts computed on these data. The psychomotor-activating effects of cocaine within each experimental group (secondary outcomes) were all unambiguous and reported in Table 2.

**Table 1.** Statistical results from the crossed contrasts computed on data dealing with the three measures of chronic psychomotor responsiveness to cocaine.

|  | Initiation of sensitization |  | Expression of sensitization |
| --- | --- | --- | --- |
|  | AUC ground | AUC increase |  |
| Female | $d = 0.39, t_{(32)} = 1.11$<br>$p = .14$ | $d = 0.01, t_{(32)} = 0.04$<br>$p = .48$ | $d = 0.21, t_{(32)} = 0.58$<br>$p = .28$ |
| Male | $d = -0.08, t_{(32)} = -0.22$<br>$p = .41$ | $d = -0.23, t_{(32)} = -0.64$<br>$p = .26$ | $d = -0.13, t_{(32)} = -0.36$<br>$p = .36$ |

**Table 2.** Statistical results from the simple contrasts (secondary outcomes) computed on data dealing with the psychomotor-activating effects of cocaine (*vs*. saline) within each experimental group.

|  | Acute reactivity | AUC ground | AUC increase | Expression |
| --- | --- | --- | --- | --- |
| EX / Female | $d = 1.04$<br>$t_{(32)} = 2.93$ | $d = 2.24$<br>$t_{(32)} = 6.33$ | $d = 1.53$<br>$t_{(32)} = 4.32$ | $d = 2.27$<br>$t_{(32)} = 6.41$ |
| SED / Female | $d = 2.09$<br>$t_{(32)} = 5.91$ | $d = 2.79$<br>$t_{(32)} = 7.90$ | $d = 1.54$<br>$t_{(32)} = 4.37$ | $d = 2.56$<br>$t_{(32)} = 7.23$ |
| EX / Male | $d = 1.14$<br>$t_{(32)} = 3.22$ | $d = 3.32$<br>$t_{(32)} = 9.39$ | $d = 1.82$<br>$t_{(32)} = 5.14$ | $d = 3.21$<br>$t_{(32)} = 9.09$ |
| SED / Male | $d = 1.62$<br>$t_{(32)} = 4.59$ | $d = 3.21$<br>$t_{(32)} = 9.08$ | $d = 1.50$<br>$t_{(32)} = 4.23$ | $d = 3.04$<br>$t_{(32)} = 8.59$ |

All *P*-values were significant, ranging from < 0.001 to 0.003.

**Figure 4.**
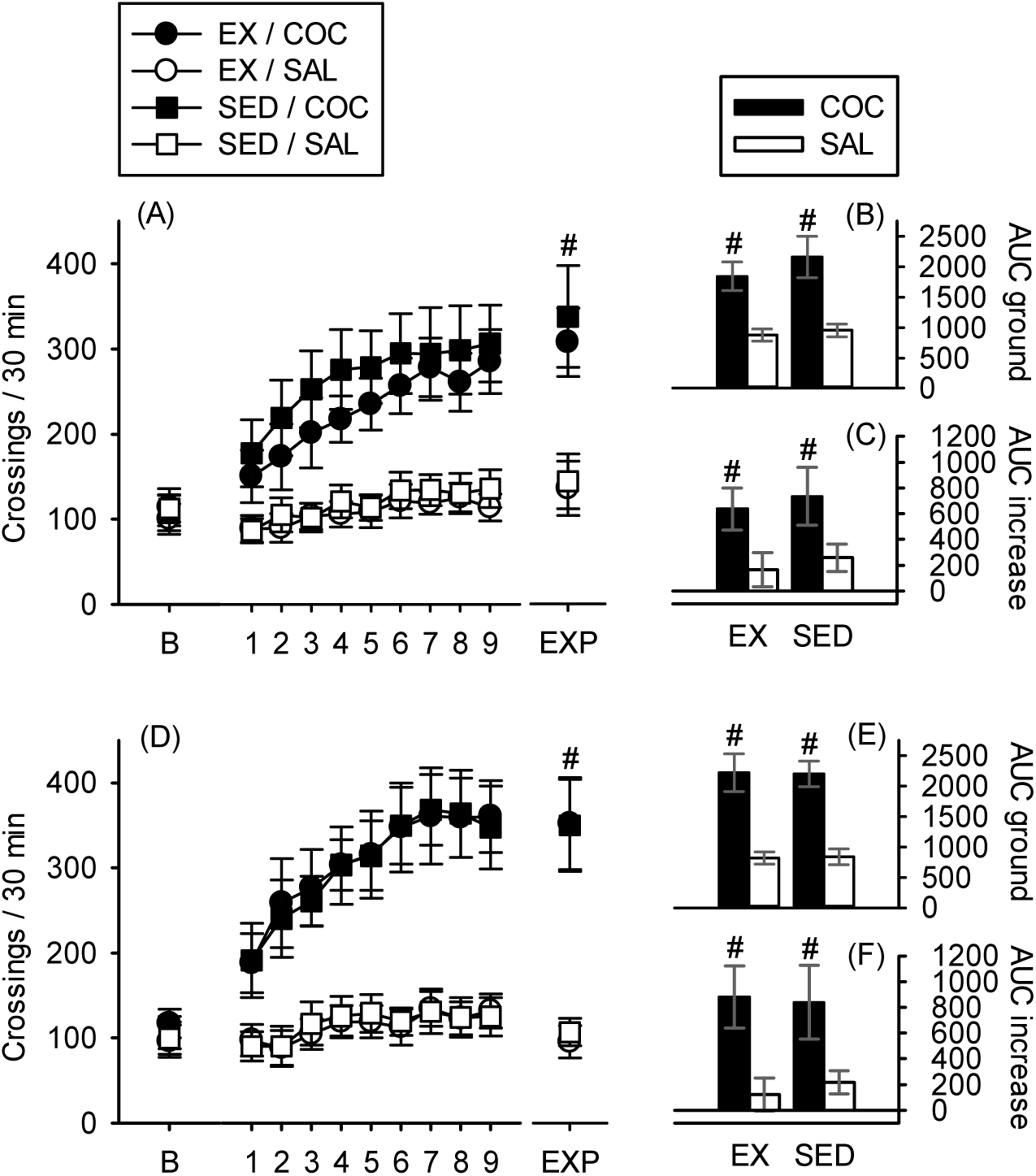
Initiation and expression of sensitization. Left panels present baseline psychomotor activity (under saline), initiation of psychomotor sensitization over 9 once-daily sessions and the long-term expression of cocaine sensitization in females (A) and males (D) born from exercised (EX) or sedentary (SED) mice before and during gestation and tested at 38-42 days of age. Right panels B (females) and E (males) show cocaine-induced psychomotor levels measured over the initiation of sensitization (overall responsiveness) scored as AUC ground. Right panels C (females) and F (males) represent cocaine-induced psychomotor sensitization levels scored as AUC increase. #: higher level than the corresponding saline group taken at a *p*-level of 0.05. Bars represent 95% CI.

## 4 Discussion

In this study we report protective-like effects of prenatal exercise (applied before and during gestation) against the acute psychomotor responsiveness to a rewarding dose of cocaine (8 mg/kg), in youth females but not in males offspring (tested at around 40 days of age). However, prenatal exercise failed to substantially influence the initiation and expression of cocaine sensitization, in either sex. Mice born from exercised and sedentary mothers displayed comparable scores or values of chronic psychomotor responsiveness to cocaine.

To the best of our knowledge, our results constitute the first report of beneficial properties conferred by maternal exercise on cocaine sensitivity in progeny, extending previous findings showing that physical activity in rodents, most often through voluntary running in a wheel, can induce protective effects on drug abuse vulnerability [3]. These results also add to a series of studies reporting that various forms of maternal aerobic exercise during gestation can positively impact neuronal and behavioural outcomes in juvenile or adult offspring, most of the studies focusing on learning and memory functions [17-20,31-33].

This set of results are in line with a plethora of research showing that brain development, which begins in utero and continues to mature until the end of adolescence [34], can be critically affected by external factors, with long-lasting consequences on health and behaviours (see [35] for a review). In particular, it is well established that early-life exposure to adverse experiences such as prenatal maternal stress (e.g., restraint) or drug of abuse administrations during pregnancy can increase the risk of cognitive and social impairments, exacerbated emotional responsiveness (e.g., anxiety or depressive-like behaviours) or drugs vulnerability in offspring [36-39]. These detrimental effects are likely due to morphological and functional changes in brain areas or circuitry involved in emotions and reward such as the dopaminergic mesolimbic pathway, hippocampus, prefrontal cortex and the hypothalamic–pituitary–adrenal axis [35,39-41]. Given accumulating evidence that exercise also acts on these systems (e.g. [42]) and that prenatal exercise can impact neuronal development, it is tempting to ascribe the exercise-induced beneficial effects to a sort of cross-tolerance phenomenon (from maternal activity to offspring cocaine effects), at least to some extent. However, the mechanisms by which maternal exercise may affect vulnerability to cocaine in female remain unknown.

Moreover, in our study, offspring were raised by their biological mothers. Therefore, one cannot rule out the possibility that changes in quality of mother-pup interactions account for the reported effects. Indeed, maternal care can be affected by environmental conditions during pregnancy such as restraint stress [43]. In this regard, further studies using cross-fostering design would be of great value to clarify the potential role of postnatal environment (maternal care) on the behavioural pattern in progeny born from exercised mothers.

The fact that the reducing effect of prenatal exercise on acute cocaine reactivity was strong in females (*d* = 0.75), but not significant and modest in males (*d* = 0.34), is in conceptual agreement with our previous study showing that the long-term protection of a 3-week regimen of wheel-running exercise during adolescence against acute and chronic psychomotor reactivity to cocaine was more pronounced in females [7]. Overall, these results also concord with a recent review of sex differences in the interaction between exercise and drugs vulnerability suggesting that females may receive more benefits from exercise than males [44]. Interestingly, Maruoka and colleagues [45] reported in C57BL/6J that environmental enrichment during gestation (comprising a running wheel) induced a (qualitative) sex-dependent effect on offspring exploratory behaviours. Maternal enrichment increased the time spent in the center of an arena (open-field test) in female only, a marker of reduced levels of anxiety-like emotional state. Since anxiolytic properties of exercise [46] have been proposed to have a mediating role in its reducing effects of drug vulnerability [3], one can thus hypothesize that the reduction of the acute psychomotor reactivity to cocaine observed in our female may rely on such mechanisms, at least in part.

Importantly, prenatal exercise did not substantially modulate (mitigate) the initiation and expression of cocaine psychomotor responsiveness, the attenuated scores of reactivity in females born from exercised mothers being limited to those dealing with acute administration. This result contrasts with the well-marked long-term protection against chronic cocaine psychomotor responsiveness previously observed in females exercised during adolescence and tested 3 weeks later as adults [7]. This set of findings indicate that, unlike adolescence, exercise before and during gestation may have a limited impact on drugs of abuse vulnerability in later life stages. In spite of being continuously accessible and voluntarily performed - thus hard to be further maximized -, one can assume that the amount of exercise during gestation was not sufficient to induce reliable and detectable long-lasting behavioural changes in our mice. Nevertheless, this explanation would conflict with the few studies describing significant changes after maternal exercise in offspring tested during late adolescence or early adulthood (from PND 40-to-60 onwards). That said, one should note that these experiments, reporting exercise-induced pro-cognitive effects, have assessed “spontaneous” exploratory behaviours, from which habituation, spatial learning or object recognition memory were inferred [18-20]. In our study, chronic cocaine administrations, mobilizing dopaminergic and glutamatergic neurotransmission within the mesolimbic system and stress-responsive functions notably (e.g., the hypothalamic-pituitary-adrenal axis) [47,48], might have been overwhelming, eventually preventing (or counteracting) the underlying mechanisms responsible for the initial protective effects seen in females to be maintained over time.

The weakness of maternal exercise effects regarding chronic psychomotor measures (*d*s varying from −0.23 to 0.39) could also rely on the length of the interval between the end of exercise (just before birth) and the psychopharmacological measures (from around 42 days of age for the initiation of sensitization and at 81 days old for its expression). Testing progeny in early adolescence (e.g., as “prepubescents”; PND 21-to-34 [49]) may provide better chances to detect exercise-induced protective effects against cocaine sensitization (chronic outcome), while remaining conceptually compatible with the literature on the ontogenesis of psychostimulants sensitization [49].

To the best of our knowledge, this study constitutes the first report of an attenuating effect of maternal exercise (before and during gestation) on cocaine responsiveness in offspring females. Further studies are warranted to characterize the interaction between prenatal exercise and drugs of abuse vulnerability in the progeny. In particular, evaluating such protection by using different models of addiction (e.g., self-administration) would be of great value in the understanding of the contribution of maternal exercise to the reduction of sensitivity to addictive properties of drugs at later life stages.

## Supporting information

## Acknowledgments

The first author (LFL) was under contract (doctoral fellow) with the Fund for Scientific Research – FNRS (Belgium) during the realization of the present work.

## References

[1] Hassandra, M., Goudas, M., & Theodorakis, Y. (2015). Exercise and Smoking: A Literature Overview. Health, 07(11), 1477–1491. https://doi.org/10.4236/health.2015.711162

[2] Lynch, W. J., Peterson, A. B., Sanchez, V., Abel, J., & Smith, M. A. (2013). Exercise as a Novel Treatment for Drug Addiction: A Neurobiological and Stage-Dependent Hypothesis. Neuroscience and Biobehavioral Reviews, 37(8), 1622–1644. https://doi.org/10.1016/j.neubiorev.2013.06.011

[3] Bardo, M. T., & Compton, W. M. (2015). Does physical activity protect against drug abuse vulnerability? Drug and Alcohol Dependence, 153, 3–13. https://doi.org/10.1016/j.drugalcdep.2015.05.037

[4] Renteria Diaz, L., Siontas, D., Mendoza, J., & Arvanitogiannis, A. (2013). High levels of wheel running protect against behavioral sensitization to cocaine. Behavioural Brain Research, 237, 82–85. https://doi.org/10.1016/j.bbr.2012.09.014

[5] Geuzaine, A., & Tirelli, E. (2014). Wheel-running mitigates psychomotor sensitization initiation but not post-sensitization conditioned activity and conditioned place preference induced by cocaine in mice. Behavioural Brain Research, 262, 57–67. https://doi.org/10.1016/j.bbr.2014.01.002

[6] Lespine, L.-F., & Tirelli, E. (2015). The protective effects of free wheel-running against cocaine psychomotor sensitization persist after exercise cessation in C57BL/6J mice. Neuroscience, 310, 650–664. https://doi.org/10.1016/j.neuroscience.2015.10.009

[7] Lespine, L.-F., & Tirelli, E. (2018). Evidence for a long-term protection of wheel-running exercise against cocaine psychomotor sensitization in adolescent but not in adult mice. Behavioural Brain Research, 349, 63–72. https://doi.org/10.1016/j.bbr.2018.04.054

[8] Leyton, M., & Vezina, P. (2013). Striatal ups and downs: Their roles in vulnerability to addictions in humans. Neuroscience & Biobehavioral Reviews, 37(9), 1999–2014. https://doi.org/10.1016/j.neubiorev.2013.01.018

[9] Steketee, J. D., & Kalivas, P. W. (2011). Drug Wanting: Behavioral Sensitization and Relapse to Drug-Seeking Behavior. Pharmacological Reviews, 63(2), 348–365. https://doi.org/10.1124/pr.109.001933

[10] Doremus-Fitzwater, T. L., Varlinskaya, E. I., & Spear, L. P. (2010). Motivational systems in adolescence: Possible implications for age differences in substance abuse and other risk-taking behaviors. Brain and Cognition, 72(1), 114. https://doi.org/10.1016/j.bandc.2009.08.008

[11] Wahlstrom, D., White, T., & Luciana, M. (2010). Neurobehavioral Evidence for Changes in Dopamine System Activity During Adolescence. Neuroscience and Biobehavioral Reviews, 34(5), 631–648. https://doi.org/10.1016/j.neubiorev.2009.12.007

[12] Committee Opinion No. 650: Physical Activity and Exercise During Pregnancy and the Postpartum Period. (2015). Obstetrics & Gynecology, 126(6), e135–e142. https://doi.org/10.1097/AOG.0000000000001214

[13] Barakat, R., Perales, M., Garatachea, N., Ruiz, J. R., & Lucia, A. (2015). Exercise during pregnancy. A narrative review asking: what do we know? British Journal of Sports Medicine, 49(21), 1377. https://doi.org/10.1136/bjsports-2015-094756

[14] Riemann, M. K., & Hansen, I.-L. K. (2000). Effects on the foetus of exercise in pregnancy. Scandinavian Journal of Medicine & Science in Sports, 10(1), 12–19. https://doi.org/10.1034/j.1600-0838.2000.010001012.x

[15] Mudd, L. M., Owe, K. M., Mottola, M. F., & Pivarnik, J. M. (2013). Health Benefits of Physical Activity during Pregnancy: An International Perspective. Medicine & Science in Sports & Exercise, 45(2), 268–277. https://doi.org/10.1249/MSS.0b013e31826cebcb

[16] Moyer, C., Reoyo, O. R., & May, L. (2016). The Influence of Prenatal Exercise on Offspring Health: A Review. Clinical Medicine Insights: Women’s Health, 9, CMWH.S34670. https://doi.org/10.4137/CMWH.S34670

[17] Aksu, I., Baykara, B., Ozbal, S., Cetin, F., Sisman, A. R., Dayi, A., … Uysal, N. (2012). Maternal treadmill exercise during pregnancy decreases anxiety and increases prefrontal cortex VEGF and BDNF levels of rat pups in early and late periods of life. Neuroscience Letters, 516(2), 221–225. https://doi.org/10.1016/j.neulet.2012.03.091

[18] Gomes da Silva, S., Unsain, N., Mascó, D. H., Toscano-Silva, M., de Amorim, H. A., Silva Araújo, B. H., … Arida, R. M. (2012). Early exercise promotes positive hippocampal plasticity and improves spatial memory in the adult life of rats. Hippocampus, 22(2), 347–358. https://doi.org/10.1002/hipo.20903

[19] Parnpiansil, P., Jutapakdeegul, N., Chentanez, T., & Kotchabhakdi, N. (2003). Exercise during pregnancy increases hippocampal brain-derived neurotrophic factor mRNA expression and spatial learning in neonatal rat pup. Neuroscience Letters, 352(1), 45–48. https://doi.org/10.1016/j.neulet.2003.08.023

[20] Robinson, A. M., & Bucci, D. J. (2014). Physical Exercise During Pregnancy Improves Object Recognition Memory in Adult Offspring. Neuroscience, 256. https://doi.org/10.1016/j.neuroscience.2013.10.012

[21] Kilkenny, C., Browne, W. J., Cuthill, I. C., Emerson, M., & Altman, D. G. (2010). Improving bioscience research reporting: The ARRIVE guidelines for reporting animal research. Journal of Pharmacology & Pharmacotherapeutics, 1(2), 94–99. https://doi.org/10.4103/0976-500X.72351

[22] Laviola, G., Macrı, S., Morley-Fletcher, S., & Adriani, W. (2003). Risk-taking behavior in adolescent mice: psychobiological determinants and early epigenetic influence. Neuroscience & Biobehavioral Reviews, 27(1–2), 19–31. https://doi.org/10.1016/S0149-7634(03)00006-X

[23] Kirk, R. E. (1995). Experimental design□: procedures for the behavioral sciences / (3rd ed.). Pacific Grove, Calif.□: Brooks/Cole.

[24] Pruessner, J. C., Kirschbaum, C., Meinlschmid, G., & Hellhammer, D. H. (2003). Two formulas for computation of the area under the curve represent measures of total hormone concentration versus time-dependent change. Psychoneuroendocrinology, 28(7), 916–931. https://doi.org/10.1016/S0306-4530(02)00108-7

[25] Cohen, J. (1988). Statistical power analysis for the behavioral sciences. Hillsdale, N.J.: L. Erlbaum Associates.

[26] Rosenthal, R., & Rosnow, R. L. (2009). Contrast analysis: focused comparisons in the analysis of variance. Cambridge: Cambridge Univ. Pr.

[27] Sawilowsky, S. S. (2009). New Effect Size Rules of Thumb. Journal of Modern Applied Statistical Methods, 8(2), 597–599. https://doi.org/10.22237/jmasm/1257035100

[28] Karasawa, K., Suwa, J., & Kimura, S. (1981). Voluntary exercise during pregnancy and lactation and its effect on lactational performance in mice. Journal of Nutritional Science and Vitaminology, 27(4), 333–339.

[29] Kelly, S. A., Hua, K., Wallace, J. N., Wells, S. E., Nehrenberg, D. L., & Pomp, D. (2015). Maternal exercise before and during pregnancy does not impact offspring exercise or body composition in mice. Journal of Negative Results in BioMedicine, 14(1). https://doi.org/10.1186/s12952-015-0032-x

[30] Zhao, Z.-J., Król, E., Moille, S., Gamo, Y., & Speakman, J. R. (2013). Limits to sustained energy intake. XV. Effects of wheel running on the energy budget during lactation. The Journal of Experimental Biology, 216(12), 2316. https://doi.org/10.1242/jeb.078402

[31] Dayi, A., Agilkaya, S., Ozbal, S., Cetin, F., Aksu, I., Gencoglu, C., … Uysal, N. (2012). Maternal Aerobic Exercise during Pregnancy Can Increase Spatial Learning by Affecting Leptin Expression on Offspring’s Early and Late Period in Life Depending on Gender. The Scientific World Journal, 2012. https://doi.org/10.1100/2012/429803

[32] Kim, H., Lee, S.-H., Kim, S.-S., Yoo, J.-H., & Kim, C.-J. (2007). The influence of maternal treadmill running during pregnancy on short-term memory and hippocampal cell survival in rat pups. International Journal of Developmental Neuroscience, 25(4), 243–249. https://doi.org/10.1016/j.ijdevneu.2007.03.003

[33] Lee, H.-H., Kim, H., Lee, J.-W., Kim, Y.-S., Yang, H.-Y., Chang, H.-K., … Kim, C.-J. (2006). Maternal swimming during pregnancy enhances short-term memory and neurogenesis in the hippocampus of rat pups. Brain and Development, 28(3), 147–154. https://doi.org/10.1016/j.braindev.2005.05.007

[34] Rice, D., & Barone, S. (2000). Critical periods of vulnerability for the developing nervous system: evidence from humans and animal models. Environmental Health Perspectives, 108(Suppl 3), 511–533.

[35] Maccari, S., Krugers, H. J., Morley-Fletcher, S., Szyf, M., & Brunton, P. J. (2014). The Consequences of Early-Life Adversity: Neurobiological, Behavioural and Epigenetic Adaptations. Journal of Neuroendocrinology, 26(10), 707–723. https://doi.org/10.1111/jne.12175

[36] Hamilton, D. A., Akers, K. G., Rice, J. P., Johnson, T. E., Candelaria-Cook, F. T., Maes, L. I., … Savage, D. D. (2010). Prenatal exposure to moderate levels of ethanol alters social behavior in adult rats: Relationship to structural plasticity and immediate early gene expression in frontal cortex. Behavioural Brain Research, 207(2), 290. https://doi.org/10.1016/j.bbr.2009.10.012

[37] Fodor, A., Tímár, J., & Zelena, D. (2014). Behavioral effects of perinatal opioid exposure. Life Sciences, 104(1), 1–8. https://doi.org/10.1016/j.lfs.2014.04.006

[38] Sanchez Vega, M. C., Chong, S., & Burne, T. H. J. (2013). Early gestational exposure to moderate concentrations of ethanol alters adult behaviour in C57BL/6J mice. Behavioural Brain Research, 252, 326–333. https://doi.org/10.1016/j.bbr.2013.06.003

[39] Hausknecht, K., Haj-Dahmane, S., & Shen, R.-Y. (2013). Prenatal Stress Exposure Increases the Excitation of Dopamine Neurons in the Ventral Tegmental Area and Alters Their Responses to Psychostimulants. Neuropsychopharmacology, 38(2), 293–301. https://doi.org/10.1038/npp.2012.168

[40] Bustamante, C., Bilbao, P., Contreras, W., Martínez, M., Mendoza, A., Reyes, Á., & Pascual, R. (2010). Effects of prenatal stress and exercise on dentate granule cells maturation and spatial memory in adolescent mice. International Journal of Developmental Neuroscience, 28(7), 605–609. https://doi.org/10.1016/j.ijdevneu.2010.07.229

[41] Bustamante, C., Henríquez, R., Medina, F., Reinoso, C., Vargas, R., & Pascual, R. (2013). Maternal exercise during pregnancy ameliorates the postnatal neuronal impairments induced by prenatal restraint stress in mice. International Journal of Developmental Neuroscience: The Official Journal of the International Society for Developmental Neuroscience, 31(4), 267–273. https://doi.org/10.1016/j.ijdevneu.2013.02.007

[42] Greenwood, B. N., Foley, T. E., Le, T. V., Strong, P. V., Loughridge, A. B., Day, H. E. W., & Fleshner, M. (2011). Long-term voluntary wheel running is rewarding and produces plasticity in the mesolimbic reward pathway. Behavioural Brain Research, 217(2), 354–362. https://doi.org/10.1016/j.bbr.2010.11.005

[43] Champagne, F. A., & Meaney, M. J. (2006). Stress During Gestation Alters Postpartum Maternal Care and the Development of the Offspring in a Rodent Model. Biological Psychiatry, 59(12), 1227–1235. https://doi.org/10.1016/j.biopsych.2005.10.016

[44] Zhou, Y., Zhao, M., Zhou, C., & Li, R. (2016). Sex differences in drug addiction and response to exercise intervention: From human to animal studies. Frontiers in Neuroendocrinology, 40, 24–41. https://doi.org/10.1016/j.yfrne.2015.07.001

[45] Maruoka, T., Kodomari, I., Yamauchi, R., Wada, E., & Wada, K. (2009). Maternal enrichment affects prenatal hippocampal proliferation and open-field behaviors in female offspring mice. Neuroscience Letters, 454(1), 28–32. https://doi.org/10.1016/j.neulet.2009.02.052

[46] Sciolino, N. R., & Holmes, P. V. (2012). Exercise offers anxiolytic potential: A role for stress and brain noradrenergic-galaninergic mechanisms. Neuroscience & Biobehavioral Reviews, 36(9), 1965–1984. https://doi.org/10.1016/j.neubiorev.2012.06.005

[47] Thomas, M. J., Kalivas, P. W., & Shaham, Y. (2009). Neuroplasticity in the mesolimbic dopamine system and cocaine addiction: Neuroplasticity and cocaine addiction. British Journal of Pharmacology, 154(2), 327–342. https://doi.org/10.1038/bjp.2008.77

[48] de Jong, I. E. M., Steenbergen, P. J., & de Kloet, E. R. (2009). Behavioral sensitization to cocaine: cooperation between glucocorticoids and epinephrine. Psychopharmacology, 204(4), 693–703. https://doi.org/10.1007/s00213-009-1498-3

[49] Tirelli, E., Laviola, G., & Adriani, W. (2003). Ontogenesis of behavioral sensitization and conditioned place preference induced by psychostimulants in laboratory rodents. Neuroscience & Biobehavioral Reviews, 27(1), 163–178. https://doi.org/10.1016/S0149-7634(03)00018-6

