## Supplementary material for "Wheel-running exercise before and during gestation against acute and sensitized cocaine psychomotor-activation in offspring"

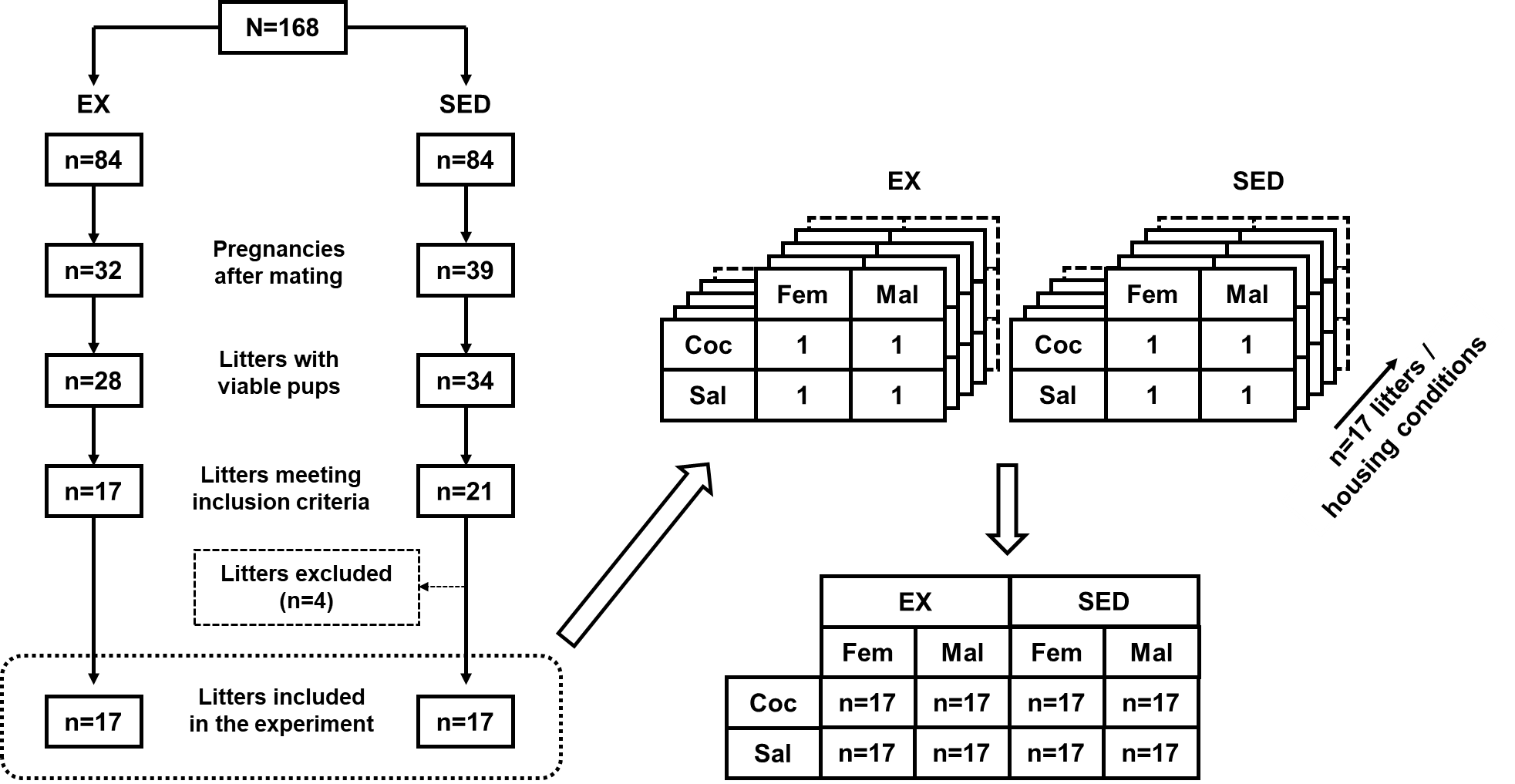

Supplement 1. A total of 168 females were used for breeding. One week after arrival, they were randomly assigned to exercise (EX, n=84) or sedentary (SED, n=84) housing. After 20 days in these conditions, they were housed with a breeder male for mating during a 12-to-60 hour-period and returned in the same pre-mating conditions (EX or SED). A total of 71 females (42.3%) showed gestation (EX, n=32; SED n=39). At birth, 9 litters were excluded due to death or not viable neonates (EX, n=4; SED, n=5). As the number of pups in C57BL/6J average 6, and because we wanted each experimental group (four groups; females and males receiving cocaine or saline) within each housing condition (EX or SED) to be represented in each litter by one mouse, only litters that comprised (at least) two females and two males were included. To employ the most homogeneous litters as possible, we also excluded litters with a number of pups ≥ 8. A total of 38 litters met these criteria (EX, n=17; SED, n=21). To obtain an equal sample size, 4 sedentary litters were excluded (randomly). Therefore, 34 litters were included in this experiment (EX, n=17; SED, n=17). Since the four possible treatments (female/cocaine, female/saline, male/cocaine, male/saline) were represented within each litter with one mouse, litter was incorporated as a random factor in a split-plot factorial design with the housing condition (2 levels: EX, SED) as a between-block factor and sex (2 levels: female, male) and drug (2 levels: cocaine, saline) as within-block factors with a total of eight experimental groups with n=17 (as independent experimental units).
