## Supplementary material for "Wheel-running exercise before and during gestation against acute and sensitized cocaine psychomotor-activation in offspring"

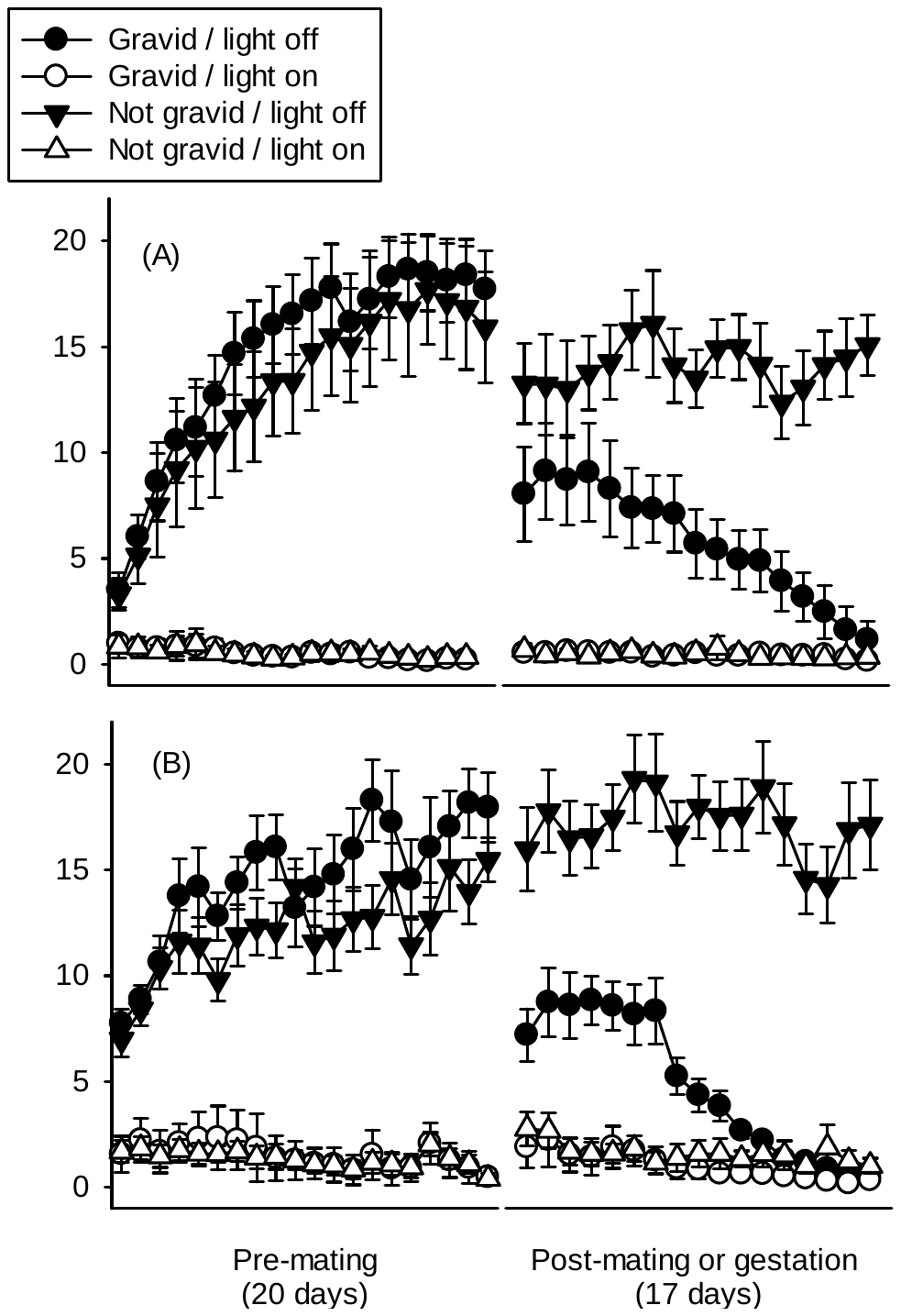

Supplement 2. Nocturnal (light off) and diurnal (light on) wheel-running activity in gravid (n=17) and not gravid (n=17) C57BL/6J (panel A; this study) and OF1 (panel B; unpublished results) females. Not gravid females were housed with a running wheel throughout the prenatal period but did not show pregnancy after the mating period. Bars represent 95% CI.
